# Myelination of major white matter tracts continues beyond childhood—combining tractography and myelin water imaging

**DOI:** 10.1101/622233

**Authors:** Tobias W. Meissner, Erhan Genç, Burkhard Mädler, Sarah Weigelt

## Abstract

Axonal myelination is a key white matter maturation process as it increases conduction velocity, synchrony, and reliability. While diffusion tensor imaging (DTI) is sensitive to myelination, it is also sensitive to unrelated microstructural properties, thus hindering straightforward interpretations. Myelin water imaging (MWI) provides a more reliable and direct *in vivo* measure of myelination. Although early histological studies show protracted myelination from childhood to adulthood, reliable tract-specific *in vivo* evidence from MWI is still lacking. Here, we combine MWI and DTI tractography to investigate myelination in middle childhood, late childhood, and adulthood in 18 major white matter tracts. In the vast majority of major white matter tracts, myelin water fraction continued to increase beyond late childhood. Our study provides first *in vivo* evidence for protracted myelination beyond late childhood.

## 1 Introduction

Initial or increasing axon myelination is a key processes of microstructural human white matter maturation as it enhances signal transmission in terms of higher transmission velocity, synchrony, and reliability (Fields, 2008). The gold standard for measuring myelin content with high accuracy are histological post-mortem studies. So far, these studies have focused on preterm and full-term neonates and young infants, finding that myelination in most parts of the brain reaches adult levels within one or two years after birth (Brody et al., 1987; Keene and Hewer, 1931; Kinney et al., 1988). However, studies that include specimens from healthy subjects between middle childhood and late adolescence are rare—maybe due no or low expectance for further myelin maturation in older years. Moreover, the few existing studies mostly focused on gray—not white—matter myelination (Abrahám et al., 2010; Benes, 1994; Benes, 1989; Flechsig, 1920; Kaes, 1907; Miller et al., 2012; Yakovlev and Lecours, 1967). They unanimously report region-specific trajectories of myelination that are prolonged beyond childhood in some areas. For white matter tracts, the same pattern was found (Yakovlev and Lecours, 1967), but evidence should be regarded as anecdotal due to the non-quantitative method of investigation and low number of investigated tracts and specimens in that age range.

The advance of diffusion tensor imaging (DTI), a magnetic resonance imaging (MRI) technique that is sensitive to myelin, non-invasive, and applicable *in vivo* lead to a large body of literature on healthy white matter maturation. Longitudinal and cross-sectional studies alike found that maturation rates peak in infancy and early childhood, but continue through childhood, adolescence, and young adulthood (e.g. Barnea-Goraly et al., 2005; Bonekamp et al., 2007; Eluvathingal et al., 2007; Lebel et al., 2008; Lebel and Beaulieu, 2011; Mukherjee et al., 2001; Schmithorst et al., 2002).

DTI achieves its sensitivity to myelin through its sensitivity to the diffusion magnitude and direction of water molecules, as new or thicker myelin reduces the inter-axonal space and thus increases the anisotropy of water diffusion (Feldman et al., 2010). However, the diffusivity of water is influenced by numerous other factors that are independent of myelin, including axonal packing density, intra-voxel orientational dispersion, membrane permeability, oligodendrocyte proliferation, and tissue water content (Jones et al., 2013; Lebel et al., 2017; Mädler et al., 2008). Thus, the interpretation of DTI parameters as markers of myelin as well as the often adapted vague term of “white matter integrity” can be problematic in studies of healthy development (Jones et al., 2013).

Recent advances in MRI have provided researchers with more direct measurements of myelin in the brain. One prominent technique is myelin water imaging (MWI). It exploits the different T2-relaxation times of water trapped between myelin sheaths and unrestricted, free intra-and extracellular water, yielding the myelin water fraction (MWF) parameter as a proxy for myelination (MacKay et al., 1994; Uddin et al., 2018). The accuracy of MWI has been validated in studies correlating histopathological myelin samples and neuroimaging-derived MWF in animal (Beaulieu et al., 1998; Gareau et al., 2000; Odrobina et al., 2005; Stanisz et al., 2004; Webb et al., 2003) and human studies (Laule et al., 2008; Laule et al., 2006).

So far, only few studies used MWI to study healthy white matter maturation. Using a variant of MWI (mcDESPOT, Deoni et al., 2008), a series of studies that investigated MWF in children up to 7 years report a steep increase of myelin from birth to age 2 and a moderate further increase thereafter. (Dean et al., 2015; Dean et al., 2014; Deoni et al., 2015; Deoni et al., 2012; Deoni et al., 2011). These findings were confirmed recently with an MR fingerprinting approach to obtain MWF data (Chen et al., 2019). A recent study that investigated nine major white matter tracts in in a cohort of twenty-three 8-13-year-olds did not find significant correlations between MWF and age (Geeraert et al., 2018). However, as histological findings suggest that myelin development might exceed late childhood in some white matter tracts and previous as tract-specific neuroimaging studies did not include adults as a reference group, conclusive evidence on myelin development beyond childhood is still lacking.

Here, our aim was to investigate myelin development between childhood and adulthood *in vivo* by combining MWI with DTI-based tractography. To this end, we measured MWF in 18 major white matter tracts in 7-8-year-old children (7-8yo), 11-12-year-old children (11-12yo), and adults (19-24yo).

## 2 Results

Eighteen children aged 7-8, 14 children aged 11-12, and 16 adults aged 19-24 were analyzed for our study. Participants completed an MRI protocol at a 3T scanner including a high-resolution T1-weighed scan, a DTI scan with 33 isotropically distributed directions, and a 3D multi-echo gradient spin echo (GRASE) MWI sequence. We reconstructed 18 major white matter tracts from DTI using probabilistic tractography (FreeSurfer TRACULA) and calculated the weighed mean myelin water fraction (MWF) for each tract in each participants’ individual native DTI space. We used independent ANOVAs for each tract to test for differences between age groups. To correct for multiple comparison, the default significance threshold of α = .05 was bonferroni-corrected to α = .05/18 = .0027

### 2.1 Myelin water fraction

Our data revealed increasing MWF with age in 13 out of 18 tracts (Figure 1, S1 Table). Effect sizes for these 13 tracts were consistently high (range of η^2^ = [.241 .704]). Increases were also evident in the other five tracts (rUNC, lILF, lCAB, rCAB, and FMIN), but did not meet our bonferroni-corrected significance threshold and featured lower effect sizes (range of η^2^ = [.143 .198], Figure 1).

**Figure 1:**
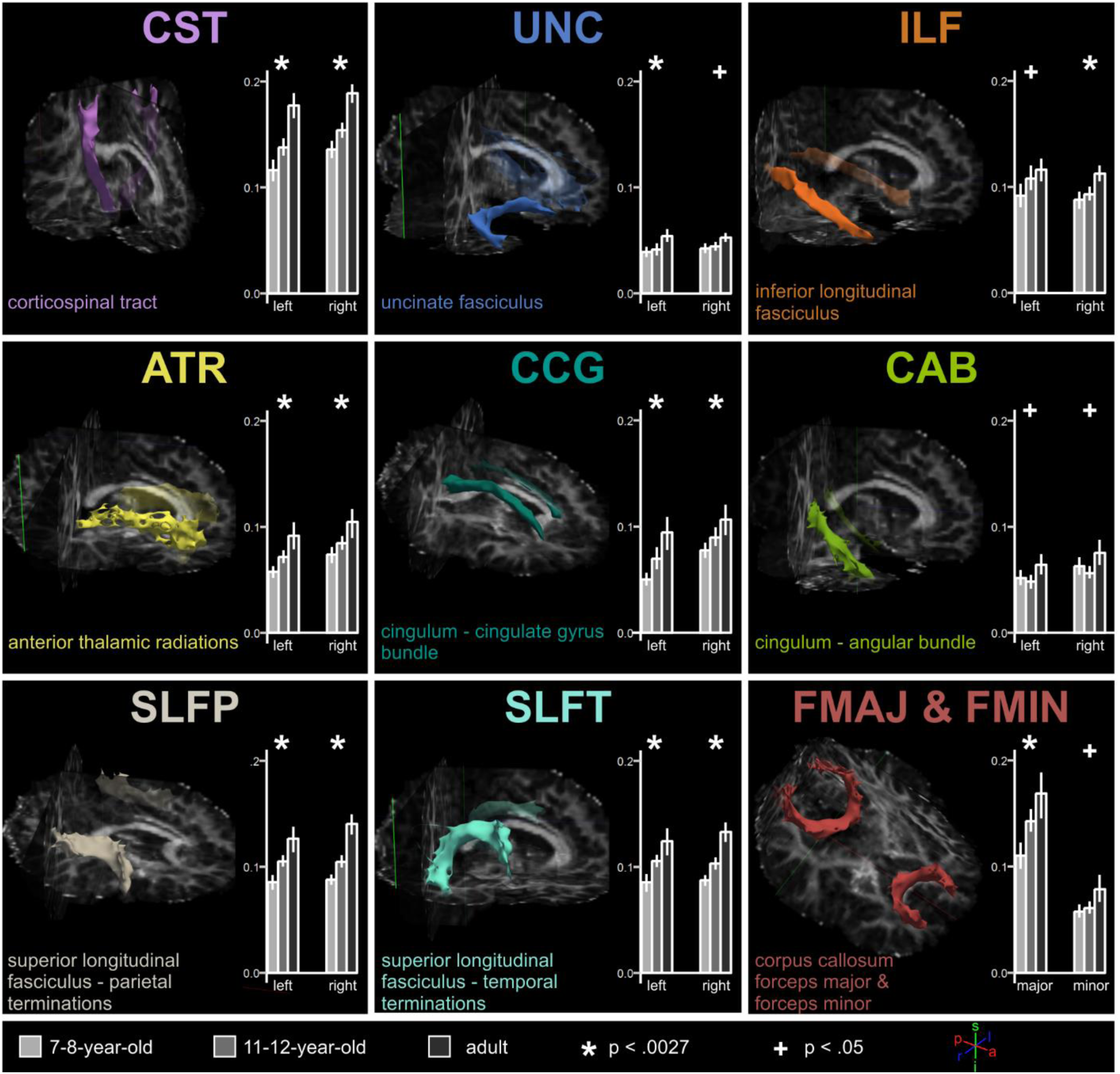
Myelin water fraction (MWF) for 18 major white matter tracts in three age groups. Light gray bars = 7-8-year-old children, medium gray bars = 11-12-year-old children, dark gray bars = adults. Error bars show 95% confidence intervals for the mean. Asterix indicate significance with Bonferroni correction (α = .0027) and plus signs without correction for multiple comparisons (α = .05), respectively. Tracts of a representative 7-8-year-old child are displayed together with one semitransparent coronal, axial, and sagittal fractional anisotropy (FA) map slice (green, blue, and red frames). All images show the brain from an anterior (a), superior (s), and right-sided (r) perspective. Within these constraints, each tract is viewed from an individual angle to emphasize its orientation in the brain. Tracts were reconstructed using the FreeSurfer TRACULA tool and a posteriori tract probability distributions were thresholded at 20% of the tract’s robust maximum probability. Isosurfaces were smoothed for visualization purposes.

In-detail, analyses of the age effects using planned comparisons *t*-tests revealed that middle and late childhood groups did not display significant differences after correction for multiple comparisons. Effect sizes in tracts that revealed significance without correction (9 out of 18 tracts), were moderate to high (range of *r* = .308 - .407, S2 Table). On the other hand, comparing children (pooled age groups) to adults yielded significant differences in all 13 tracts that yielded an age effect after correction for multiple comparisons, with consistently high effect sizes (range of *r* = [.427 .836], S2 Table). In summary, our data suggests strong protracted increases of MWF that seem to be driven by changes well beyond late childhood.

### 2.2 Supplementary analysis of diffusion parameters

In a supplementary analysis, we investigated if DTI parameters fractional anisotropy (FA), mean diffusivity (MD), radial diffusivity (RD) and axial diffusivity (AD) mirrored the observed MWI effects. In short, MD and RD decreased and mirrored myelin development in most tracts, while FA and AD matched MWF age effects in fewer tracts (S1 Text, S2 Figure, S3-S6 Tables). We demonstrate that interpreting DTI parameters as a proxy for myelination is less accurate than MWI in detecting age differences. Still, MD and RD can reproduce myelin-specific age effects in most tracts (S2 Text).

## 3 Discussion

Our study compared MWF in 18 major white matter tracts of 7-8yo, 11-12yo and young adults and found that MWF substantially increases in 13 tracts. Thus, we provide for the first time the *in vivo* evidence for a continued increase of myelin throughout childhood and adolescence.

Previous studies using MWI reported drastic increases of myelin in infancy but lower rates of development in early and middle childhood (Chen et al., 2019; Dean et al., 2015; Dean et al., 2014; Deoni et al., 2015; Deoni et al., 2012; Deoni et al., 2011) and a stagnation in middle to late childhood (Geeraert et al., 2018). However, our study paints a different picture. In contrast to earlier MWI studies, we present data from middle and late childhood in comparison to adult data. This allows for the first time the investigation of further protracted myelin development beyond late childhood. Our MWI data suggest that myelination is not limited to infancy and early childhood. Rather, it continues well beyond childhood, at least until early adolescence, and possibly until adulthood in major white matter tracts. Thus, using MWI, we support early histological reports (Benes, 1989; Yakovlev and Lecours, 1967) as well as DTI studies (e.g. Barnea-Goraly et al., 2005; Bonekamp et al., 2007; Eluvathingal et al., 2007; Lebel et al., 2008; Lebel and Beaulieu, 2011; Schmithorst et al., 2002) claiming that myelin development is protracted until adulthood.

Our in-detail planned comparisons analyses revealed that age effects were strongly driven by increasing MWF between children and adults, rather than between 7-8yo and 11-12yo—a pattern with two implications. First, myelination might increase at a very moderate pace—if at all—between early and late childhood. This implication and our data concur with a recent study that correlated MWF and age in 8-13-year-old children (Geeraert et al., 2018). While the authors found correlations of MWF with age in some tracts, multiple comparison corrections rendered these relationships as not significant, as in our comparison of 7-8yo and 11-12yo. Second, our result pattern implies that age effects in our study are driven by myelination events between late childhood (11-12yo) and adulthood. Integrating this finding with the plateau-like trajectory in early childhood studies and our lower myelination rates between middle and late childhood, this might indicate a pronounced myelin increase, possibly even a short second boost, in adolescence. Changes of that magnitude might give rise to—or might stem from—changed cortical functionality during that time (Fields, 2015).

Our study sheds first light onto examining myelination development *in vivo* throughout childhood using MWI. Two strands of future research are needed to corroborate our findings and unravel the exact trajectory of myelin development beyond early childhood. First, there is a strong need for longitudinal MWI studies that pinpoint myelin growth from early childhood to adulthood with relatively high power and precision, e.g. in 2-year intervals. At the same time, the retest-reliability of current MWI techniques should be evaluated in pediatric participants to ensure meaningful inference in longitudinal settings. Second, we are in strong need for post-mortem histochemical studies with specimens from neurologically healthy children and adolescents that operate on white matter in a tract-specific and quantitative fashion to validate MWI. These future directions could enable practitioners to use MWI as a reliable clinical marker for myelin-related conditions or symptoms in childhood and enable researchers to investigate myelin-related brain-behavior associations in periods of marked behavioral changes, such as adolescence.

## 4 Methods

### 4.1 Participants

We analyzed data of 48 participants for this study (18 children aged 7-8, *M* = 7.56, *SD* = 0.51; 7 female; 14 children aged 11-12, *M* = 11.21, *SD* = 0.43; 9 female; 16 adults aged 19-24, *M* = 20.69, *SD* = 1.14; 7 female). All participants were healthy. Participants took part in a larger project with multiple MRI protocols. Details on health exclusion criteria, subject recruitment and compensation are reported elsewhere (Meissner et al., 2019). The original sample included two additional 7-8yo that were excluded due to severely impaired data quality in the DTI scan or due to subject-induced scan session termination before myelin water imaging, respectively.

### 4.2 Neuroimaging

All MRI was conducted at the Neuroimaging Centre of the Research Department of Neuroscience at Ruhr University Bochum’s teaching hospital Bergmannsheil on a 3.0T Archieva scanner (Philips, Amsterdam, The Netherlands) using a 32-channel head coil. Children were accompanied by one of the experimenters in the scanner room throughout the entire procedure and— given no prior MRI experience—were accustomed to the scanning environment in a custom-built mock scanner at least one day prior to scanning. All participants viewed episodes of a children’s TV program during non-functional MRI scans.

#### 4.2.1 High-resolution anatomical imaging

To co-register DTI and MWI as well as for gray-white matter segmentation and cortical parcellation (a prerequisite for later TRACULA tractography), we acquired a T1-weighted high-resolution anatomical scan of the whole head (MP-RAGE, TR = 8.10 s, TE = 3.72 ms, flip angle = 8°, 220 slices, matrix size = 240 × 240, voxel size = 1 mm × 1 mm × 1 mm). Brains were extracted from anatomical images using FSL BET (Smith, 2002).

#### 4.2.2 Diffusion weighted imaging

For probabilistic tractography, a diffusion-weighted single-shot spin-echo EPI sequence along 33 isotropically distributed directions was obtained (b = 1000 s/mm^2^, TR = 7234 ms, TE = 89 ms, flip angle = 90°, 60 slices, matrix size = 128 × 128, voxel size = 2 × 2 × 2mm) and preceded by one reference image without diffusion weighting (b = 0 s/mm^2^). We used FreeSurfer’s (RRID: SCR_001847) TRACULA tool (TRActs Constrained by UnderLying Anatomy, Yendiki et al., 2011) with default settings to delineate 18 major white matter tracts. Tracts included the bilateral components of the corticospinal tract (CST), uncinate fasciculus (UNC), inferior longitudinal fasciculus (ILF), anterior thalamic radiations (ATR), cingulate gyrus bundle of the cingulum (CCG), angular bundle of the cingulum (CAB), parietal terminations of the superior longitudinal fasciculus (SLFP), temporal terminations of the superior longitudinal fasciculus (SLFT), as well as the forceps major (FMAJ) and the forceps minor (FMIN) of the corpus callosum (Figure 1). Data preprocessing included eddy current correction, head motion correction, and brain extraction. A ball-and-stick model was fit to each voxel using FSL BEDPOSTX (Behrens et al., 2007; Behrens et al., 2003). For each tract, TRACULA first estimates the *a priori* probability by combining atlas data (of manually labeled tracts and anatomical segmentations in training subjects) and the subject’s automatically reconstructed and segmented anatomical T1-weighed images (obtained by FreeSurfer’s recon-all tool, Dale et al., 1999; Fischl et al., 2004; Fischl et al., 2002; Fischl et al., 1999; Ségonne et al., 2004). Then, TRACULA estimates an *a posteriori* probability distribution based on the *a priori* probability and the ball-and-stick model. The recon-all tool was shown to produce valid results for children (Ghosh et al., 2010) and the TRACULA tool does not rely on perfect alignment of individual subject data to the atlas space (Yendiki et al., 2011). Thus, we follow the developer’s assessment and judge TRACULA as a valid tool for all age groups in our study (Yendiki et al., 2014). For details on TRACULA, please refer to Yendiki et al. (2011) and the respective online documentation (https://surfer.nmr.mgh.harvard.edu/fswiki/Tracula).

To remove spurious connections indicated by low-probability voxels, we thresholded each tract’s *a posteriori* probability distribution map at 20% of its robust maximum (i.e. 99th percentile) probability.

#### 4.2.3 Myelin water imaging

For *in vivo* myelin examination, a 3D multi-echo gradient spin echo (GRASE) sequence with refocusing sweep angle was acquired (TR = 800 ms; TE = 10 - 320 ms, 32 echoes in steps of 10 ms, partial Fourier acquisition in both phase encoding directions, parallel imaging SENSE = 2.0, flip angle = 90°, 60 slices, matrix size = 112 × 112, voxel size = 2 × 2 × 2 mm, acquisition duration = 7.25 min). Parameter maps estimating the myelin water fraction (MWF, i.e. fraction of water molecules located between myelin sheath layers, MacKay et al., 1994), were created as described elsewhere (Ocklenburg et al., 2018; for details, see Prasloski et al., 2012). MWF maps were registered to native DTI space in two steps via the anatomical T1 space using sinc interpolation in FSL FLIRT (FMRIB’s Linear Image Registration Tool, Greve and Fischl, 2009; Jenkinson et al., 2002; Jenkinson and Smith, 2001). We calculated weighted mean MWF values for each tract that considered each voxel’s MWF value and tract probability (number of samples crossing through a voxel in relation to the total number of successful samples).

### 4.3 Neuroimaging data quality control

We screened raw and preprocessed 4D DTI data for visible artefacts and excluded one 7-8yo subject from subsequent analyses (see 4.1, Participants). Further, to control for possible age group differences in DTI data quality, we analyzed four data quality measures obtained by the TRACULA preprocessing tools (Yendiki et al., 2014). To capture global, slow between-volume motion, we compared volume-to-volume translation and rotation parameters between age groups. None of the parameters revealed between-group differences (S1 Figure; rotation: *F*(2,45) = 2.03, *p* = .143, η^2^ = .083 translation: *F*(2,45) = 2.45, *p* = .098, η^2^ = .098). To capture the effect of rapid within-volume motion (note: TR = 7234 ms), we compared slice signal dropout (Benner et al., 2011) between age groups. Only four 7-8yo, two 11-12yo, and no adults displayed slices with signal dropout (S1 Figure). In participants with signal dropout, the portion of dropout slices was 0.21 % at maximum. Due to the low number of participants with signal dropout, and no signal dropout in the adult group (i.e. no variance), traditional ANOVAs for percentage of dropout slices or severity of dropout scores between age groups lack validity. As an alternative, we employed Fisher’s exact test and found that the number of participants with any signal-dropout did not differ between age groups (χ^2^(2) = 3.88, *p* = .138).

As the 3D signal acquisition method of the GRASE sequence is not volume-based, affine registration matrices and corresponding motion estimates, as for fMRI or DTI, cannot be computed for 3D ME-GRASE data. However, we visually screened all raw GRASE images as well as MWF maps for motion artefacts but found none.

### 4.4 Experimental design and statistical analysis

Our study investigated the effect of the between-subject factor age group (with three levels) on the outcome variable MWF for 18 major white matter tracts. We used independent ANOVAs for each tract to test for differences between age groups. To correct for multiple comparison, the default significance threshold of α = .05 was bonferroni-corrected to α = .05/18 = .0027. To improve the usability of our results for colleagues, whose research interest focuses on one or a few tracts only, we also report age group effects that reached the uncorrected significance threshold of α = .05 in a second step. Statistical data analysis was performed using R (version 3.2.2, RRID: SCR_001905, R Core Team, 2015) in RStudio (version 0.99.491; RRID: SCR_000432).

## Supporting information

Combined Supplementary Material

## Acknowledgements

We acknowledge the support of the Neuroimaging Centre of the Research Department of Neuroscience at Ruhr University Bochum’s teaching hospital Bergmannsheil. We thank all participants and their parents for participating in this study.

## Author contributions

Conceptualization: TWM; Methodology: EG, BM; Software: BM; Formal Analysis: TWM, EG; Investigation: TWM; Resources: SW, EG; Data Curation: TWM; Writing—Original Draft: TWM; Writing—Review and Editing: TWM, SW, EG, BM; Visualization: TWM; Supervision: SW; Project Administration: TWM; Funding Acquisition: SW, TWM

## Data and code availability statement

All code used for data analysis (except for the MWF parameter map generating algorithm), as well as anonymized raw data are publicly available at the Open Science Framework (https://osf.io/v9tkz/, https://doi.org/10.17605/osf.io/v9tkz).

## Ethics statement

The Ruhr University Bochum Faculty of Psychology ethics board approved the study (proposal no. 280). All participants as well as children’s parents gave informed written consent to participate voluntarily.

## References

Abrahám, H., Vincze, A., Jewgenow, I., Veszprémi, B., Kravják, A., Gömöri, E., Seress, L., 2010. Myelination in the human hippocampal formation from midgestation to adulthood. Int J Dev Neurosci 28 (5), 401–410. 10.1016/j.ijdevneu.2010.03.004.

Barnea-Goraly, N., Menon, V., Eckert, M., Tamm, L., Bammer, R., Karchemskiy, A., Dant, C.C., Reiss, A.L., 2005. White matter development during childhood and adolescence: a cross-sectional diffusion-tensor imaging study. Cereb Cortex 15 (12), 1848–1854. 10.1093/cercor/bhi062.

Beaulieu, C., Fenrich, F.R., Allen, P.S., 1998. Multicomponent water proton transverse relaxation and T2-discriminated water diffusion in myelinated and nonmyelinated nerve. Magnetic resonance imaging 16 (10), 1201–1210. 10.1016/S0730-725X(98)00151-9.

Behrens, T.E.J., Berg, H.J., Jbabdi, S., Rushworth, M.F.S., Woolrich, M.W., 2007. Probabilistic diffusion tractography with multiple fibre orientations: What can we gain? Neuroimage 34 (1), 144–155. 10.1016/j.neuroimage.2006.09.018.

Behrens, T.E.J., Woolrich, M.W., Jenkinson, M., Johansen-Berg, H., Nunes, R.G., Clare, S., Matthews, P.M., Brady, J.M., Smith, S.M., 2003. Characterization and propagation of uncertainty in diffusion-weighted MR imaging. Magn Reson Med 50 (5), 1077–1088. 10.1002/mrm.10609.

Benes, F.M., 1989. Myelination of Cortical-hippocampal Relays During Late Adolescence. Schizophrenia Bulletin 15 (4), 585–593. 10.1093/schbul/15.4.585.

Benes, F.M., 1994. Myelination of a Key Relay Zone in the Hippocampal Formation Occurs in the Human Brain During Childhood, Adolescence, and Adulthood. Arch Gen Psychiatry 51 (6), 477. 10.1001/archpsyc.1994.03950060041004.

Benner, T., van der Kouwe, A.J.W., Sorensen, A.G., 2011. Diffusion imaging with prospective motion correction and reacquisition. Magn Reson Med 66 (1), 154–167. 10.1002/mrm.22837.

Bonekamp, D., Nagae, L.M., Degaonkar, M., Matson, M., Abdalla, W.M.A., Barker, P.B., Mori, S., Horská, A., 2007. Diffusion tensor imaging in children and adolescents: reproducibility, hemispheric, and age-related differences. Neuroimage 34 (2), 733–742. 10.1016/j.neuroimage.2006.09.020.

Brody, B.A., Kinney, H.C., Kloman, A.S., Gilles, F.H., 1987. Sequence of central nervous system myelination in human infancy. I. An autopsy study of myelination. Journal of Neuropathology and Experimental Neurology 46 (3), 283–301.

Chen, Y., Chen, M.-H., Baluyot, K.R., Potts, T.M., Jimenez, J., Lin, W., 2019. MR fingerprinting enables quantitative measures of brain tissue relaxation times and myelin water fraction in the first five years of life. Neuroimage 186, 782–793. 10.1016/j.neuroimage.2018.11.038.

Dale, A.M., Fischl, B., Sereno, M.I., 1999. Cortical surface-based analysis. I. Segmentation and surface reconstruction. Neuroimage 9 (2), 179–194. 10.1006/nimg.1998.0395.

Dean, D.C., O’Muircheartaigh, J., Dirks, H., Waskiewicz, N., Lehman, K., Walker, L., Han, M., Deoni, S.C.L., 2014. Modeling healthy male white matter and myelin development: 3 through 60months of age. Neuroimage 84, 742–752. 10.1016/j.neuroimage.2013.09.058.

Dean, D.C., O’Muircheartaigh, J., Dirks, H., Waskiewicz, N., Walker, L., Doernberg, E., Piryatinsky, I., Deoni, S.C.L., 2015. Characterizing longitudinal white matter development during early childhood. Brain Struct Funct 220 (4), 1921–1933. 10.1007/s00429-014-0763-3.

Deoni, S.C.L., Dean, D.C., O’Muircheartaigh, J., Dirks, H., Jerskey, B.A., 2012. Investigating white matter development in infancy and early childhood using myelin water faction and relaxation time mapping. Neuroimage 63 (3), 1038–1053. 10.1016/j.neuroimage.2012.07.037.

Deoni, S.C.L., Dean, D.C., Remer, J., Dirks, H., O’Muircheartaigh, J., 2015. Cortical maturation and myelination in healthy toddlers and young children. Neuroimage 115, 147–161. 10.1016/j.neuroimage.2015.04.058.

Deoni, S.C.L., Mercure, E., Blasi, A., Gasston, D., Thomson, A., Johnson, M., Williams, S.C.R., Murphy, D.G.M., 2011. Mapping infant brain myelination with magnetic resonance imaging. J Neurosci 31 (2), 784–791. 10.1523/JNEUROSCI.2106-10.2011.

Deoni, S.C.L., Rutt, B.K., Arun, T., Pierpaoli, C., Jones, D.K., 2008. Gleaning multicomponent T1 and T2 information from steady-state imaging data. Magn Reson Med 60 (6), 1372–1387. 10.1002/mrm.21704.

Eluvathingal, T.J., Hasan, K.M., Kramer, L., Fletcher, J.M., Ewing-Cobbs, L., 2007. Quantitative diffusion tensor tractography of association and projection fibers in normally developing children and adolescents. Cereb Cortex 17 (12), 2760–2768. 10.1093/cercor/bhm003.

Feldman, H.M., Yeatman, J.D., Lee, E.S., Barde, L.H.F., Gaman-Bean, S., 2010. Diffusion tensor imaging: a review for pediatric researchers and clinicians. J Dev Behav Pediatr 31 (4), 346–356. 10.1097/DBP.0b013e3181dcaa8b.

Fields, R.D., 2008. White matter in learning, cognition and psychiatric disorders. Trends Neurosci 31 (7), 361–370. 10.1016/j.tins.2008.04.001.

Fields, R.D., 2015. A new mechanism of nervous system plasticity: activity-dependent myelination. Nat Rev Neurosci 16 (12), 756–767. 10.1038/nrn4023.

Fischl, B., Salat, D.H., Busa, E., Albert, M., Dieterich, M., Haselgrove, C., van der Kouwe, A., Killiany, R., Kennedy, D., Klaveness, S., Montillo, A., Makris, N., Rosen, B., Dale, A.M., 2002. Whole Brain Segmentation. Neuron 33 (3), 341–355. 10.1016/S0896-6273(02)00569-X.

Fischl, B., Sereno, M.I., Dale, A.M., 1999. Cortical surface-based analysis. II: Inflation, flattening, and a surface-based coordinate system. Neuroimage 9 (2), 195–207. 10.1006/nimg.1998.0396.

Fischl, B., van der Kouwe, A., Destrieux, C., Halgren, E., Ségonne, F., Salat, D.H., Busa, E., Seidman, L.J., Goldstein, J., Kennedy, D., Caviness, V., Makris, N., Rosen, B., Dale, A.M., 2004. Automatically parcellating the human cerebral cortex. Cereb Cortex 14 (1), 11–22.

Flechsig, P., 1920. Anatomie des menschlichen Gehirns und Rückenmarks auf myelogenetischer Grundlage. Thieme, Leipzig.

Gareau, P.J., Rutt, B.K., Karlik, S.J., Mitchell, J.R., 2000. Magnetization transfer and multicomponent T2 relaxation measurements with histopathologic correlation in an experimental model of MS. J Magn Reson Imaging 11 (6), 586–595.

Geeraert, B.L., Lebel, R.M., Mah, A.C., Deoni, S.C., Alsop, D.C., Varma, G., Lebel, C., 2018. A comparison of inhomogeneous magnetization transfer, myelin volume fraction, and diffusion tensor imaging measures in healthy children. Neuroimage 182, 343–350. 10.1016/j.neuroimage.2017.09.019.

Ghosh, S.S., Kakunoori, S., Augustinack, J., Nieto-Castanon, A., Kovelman, I., Gaab, N., Christodoulou, J.A., Triantafyllou, C., Gabrieli, J.D.E., Fischl, B., 2010. Evaluating the validity of volume-based and surface-based brain image registration for developmental cognitive neuroscience studies in children 4 to 11 years of age. Neuroimage 53 (1), 85–93. 10.1016/j.neuroimage.2010.05.075.

Greve, D.N., Fischl, B., 2009. Accurate and robust brain image alignment using boundary-based registration. Neuroimage 48 (1), 63–72. 10.1016/j.neuroimage.2009.06.060.

Jenkinson, M., Bannister, P., Brady, M., Smith, S., 2002. Improved Optimization for the Robust and Accurate Linear Registration and Motion Correction of Brain Images. Neuroimage 17 (2), 825–841. 10.1006/nimg.2002.1132.

Jenkinson, M., Smith, S., 2001. A global optimisation method for robust affine registration of brain images. Medical Image Analysis 5 (2), 143–156. 10.1016/S1361-8415(01)00036-6.

Jones, D.K., Knösche, T.R., Turner, R., 2013. White matter integrity, fiber count, and other fallacies: the do’s and don’ts of diffusion MRI. Neuroimage 73, 239–254. 10.1016/j.neuroimage.2012.06.081.

Kaes, T., 1907. Die Grosshirnrinde des Menschen in ihren Massen und ihrem Fasergehalt. Fischer, Jena.

Keene, M.F.L., Hewer, E.E., 1931. Some Observations on Myelination in the Human Central Nervous System. J Anatomy 66 (Pt 1), 1–13.

Kinney, H.C., Brody, B.A., Kloman, A.S., Gilles, F.H., 1988. Sequence of central nervous system myelination in human infancy. II. Patterns of myelination in autopsied infants. Journal of Neuropathology and Experimental Neurology 47 (3), 217–234.

Laule, C., Kozlowski, P., Leung, E., Li, D.K.B., MacKay, A.L., Moore, G.R.W., 2008. Myelin water imaging of multiple sclerosis at 7 T: correlations with histopathology. Neuroimage 40 (4), 1575–1580. 10.1016/j.neuroimage.2007.12.008.

Laule, C., Leung, E., Lis, D.K.B., Traboulsee, A.L., Paty, D.W., MacKay, A.L., Moore, G.R.W., 2006. Myelin water imaging in multiple sclerosis: quantitative correlations with histopathology. Mult Scler 12 (6), 747–753. 10.1177/1352458506070928.

Lebel, C., Beaulieu, C., 2011. Longitudinal Development of Human Brain Wiring Continues from Childhood into Adulthood. J Neurosci 31 (30), 10937–10947. 10.1523/JNEUROSCI.5302-10.2011.

Lebel, C., Treit, S., Beaulieu, C., 2017. A review of diffusion MRI of typical white matter development from early childhood to young adulthood. NMR in biomedicine. 10.1002/nbm.3778.

Lebel, C., Walker, L., Leemans, A., Phillips, L., Beaulieu, C., 2008. Microstructural maturation of the human brain from childhood to adulthood. Neuroimage 40 (3), 1044–1055. 10.1016/j.neuroimage.2007.12.053.

MacKay, A., Whittall, K., Adler, J., Li, D., Paty, D., Graeb, D., 1994. In vivo visualization of myelin water in brain by magnetic resonance. Magn Reson Med 31 (6), 673–677.

Mädler, B., Drabycz, S.A., Kolind, S.H., Whittall, K.P., MacKay, A.L., 2008. Is diffusion anisotropy an accurate monitor of myelination? Correlation of multicomponent T2 relaxation and diffusion tensor anisotropy in human brain. Magnetic resonance imaging 26 (7), 874–888. 10.1016/j.mri.2008.01.047.

Meissner, T.W., Nordt, M., Weigelt, S., 2019. Prolonged functional development of the parahippocampal place area and occipital place area. Neuroimage 191, 104–115. 10.1016/j.neuroimage.2019.02.025.

Miller, D.J., Duka, T., Stimpson, C.D., Schapiro, S.J., Baze, W.B., McArthur, M.J., Fobbs, A.J., Sousa, A.M.M., Sestan, N., Wildman, D.E., Lipovich, L., Kuzawa, C.W., Hof, P.R., Sherwood, C.C., 2012. Prolonged myelination in human neocortical evolution. Proc Natl Acad Sci USA 109 (41), 16480–16485. 10.1073/pnas.1117943109.

Mukherjee, P., Miller, J.H., Shimony, J.S., Conturo, T.E., Lee, B.C., Almli, C.R., McKinstry, R.C., 2001. Normal brain maturation during childhood: developmental trends characterized with diffusion-tensor MR imaging. Radiology 221 (2), 349–358. 10.1148/radiol.2212001702.

Ocklenburg, S., Anderson, C., Gerding, W.M., Fraenz, C., Schlüter, C., Friedrich, P., Raane, M., Mädler, B., Schlaffke, L., Arning, L., Epplen, J.T., Güntürkün, O., Beste, C., Genç, E., 2018. Myelin Water Fraction Imaging Reveals Hemispheric Asymmetries in Human White Matter That Are Associated with Genetic Variation in PLP1. Molecular neurobiology. 10.1007/s12035-018-1351-y.

Odrobina, E.E., Lam, T.Y.J., Pun, T., Midha, R., Stanisz, G.J., 2005. MR properties of excised neural tissue following experimentally induced demyelination. NMR in biomedicine 18 (5), 277–284. 10.1002/nbm.951.

Prasloski, T., Rauscher, A., MacKay, A.L., Hodgson, M., Vavasour, I.M., Laule, C., Mädler, B., 2012. Rapid whole cerebrum myelin water imaging using a 3D GRASE sequence. Neuroimage 63 (1), 533–539. 10.1016/j.neuroimage.2012.06.064.

R Core Team, 2015. R: A Language and Environment for Statistical Computing, Vienna, Austria. http://www.R-project.org/.

Schmithorst, V.J., Wilke, M., Dardzinski, B.J., Holland, S.K., 2002. Correlation of white matter diffusivity and anisotropy with age during childhood and adolescence: a cross-sectional diffusiontensor MR imaging study. Radiology 222 (1), 212–218. 10.1148/radiol.2221010626.

Ségonne, F., Dale, A.M., Busa, E., Glessner, M., Salat, D., Hahn, H.K., Fischl, B., 2004. A hybrid approach to the skull stripping problem in MRI. Neuroimage 22 (3), 1060–1075. 10.1016/j.neuroimage.2004.03.032.

Smith, S.M., 2002. Fast robust automated brain extraction. Hum Brain Mapp 17 (3), 143–155. 10.1002/hbm.10062.

Stanisz, G.J., Webb, S., Munro, C.A., Pun, T., Midha, R., 2004. MR properties of excised neural tissue following experimentally induced inflammation. Magn Reson Med 51 (3), 473–479. 10.1002/mrm.20008.

Uddin, M.N., Figley, T.D., Marrie, R.A., Figley, C.R., 2018. Can T1 w/T2 w ratio be used as a myelin-specific measure in subcortical structures? Comparisons between FSE-based T1 w/T2 w ratios, GRASE-based T1 w/T2 w ratios and multi-echo GRASE-based myelin water fractions. NMR in biomedicine 31 (3). 10.1002/nbm.3868.

Webb, S., Munro, C.A., Midha, R., Stanisz, G.J., 2003. Is multicomponent T2 a good measure of myelin content in peripheral nerve? Magn Reson Med 49 (4), 638–645. 10.1002/mrm.10411.

Yakovlev, P.I., Lecours, A.-R., 1967. The myelogenetic cycles of regional maturation of the brain, in: Minkowski, A. (Ed.), Regional Development of the Brain Early in Life. Blackwell Scientific Publications Inc., Boston, MA, pp. 3–70.

Yendiki, A., Koldewyn, K., Kakunoori, S., Kanwisher, N., Fischl, B., 2014. Spurious group differences due to head motion in a diffusion MRI study. Neuroimage 88, 79–90. 10.1016/j.neuroimage.2013.11.027.

Yendiki, A., Panneck, P., Srinivasan, P., Stevens, A., Zöllei, L., Augustinack, J., Wang, R., Salat, D., Ehrlich, S., Behrens, T., Jbabdi, S., Gollub, R., Fischl, B., 2011. Automated probabilistic reconstruction of white-matter pathways in health and disease using an atlas of the underlying anatomy. Front Neuroinform 5, 23. 10.3389/fninf.2011.00023.

