## Supplementary material for "Myelination of major white matter tracts continues beyond childhood—combining tractography and myelin water imaging": Combined Supplementary Material

### Figures

#### S1 Figure

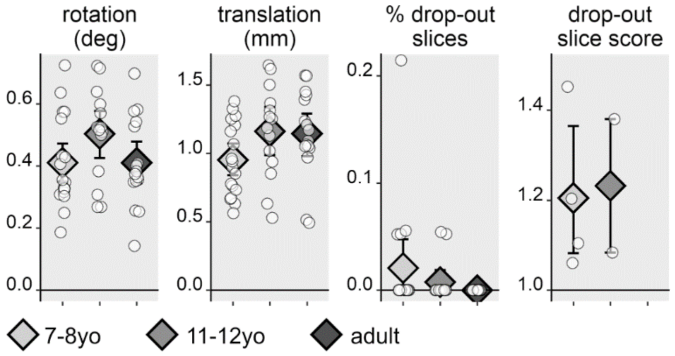

***S1 Figure**: Between-group data quality comparison after matching for the percentage of drop-out slices. Light gray = 7-8-year-old children, medium gray = 11-12-year-old children, dark gray = adults. White circles = individual data points. Gray diamonds = group mean. Error bars show 95% confidence intervals for the mean.*

### S2 Figure

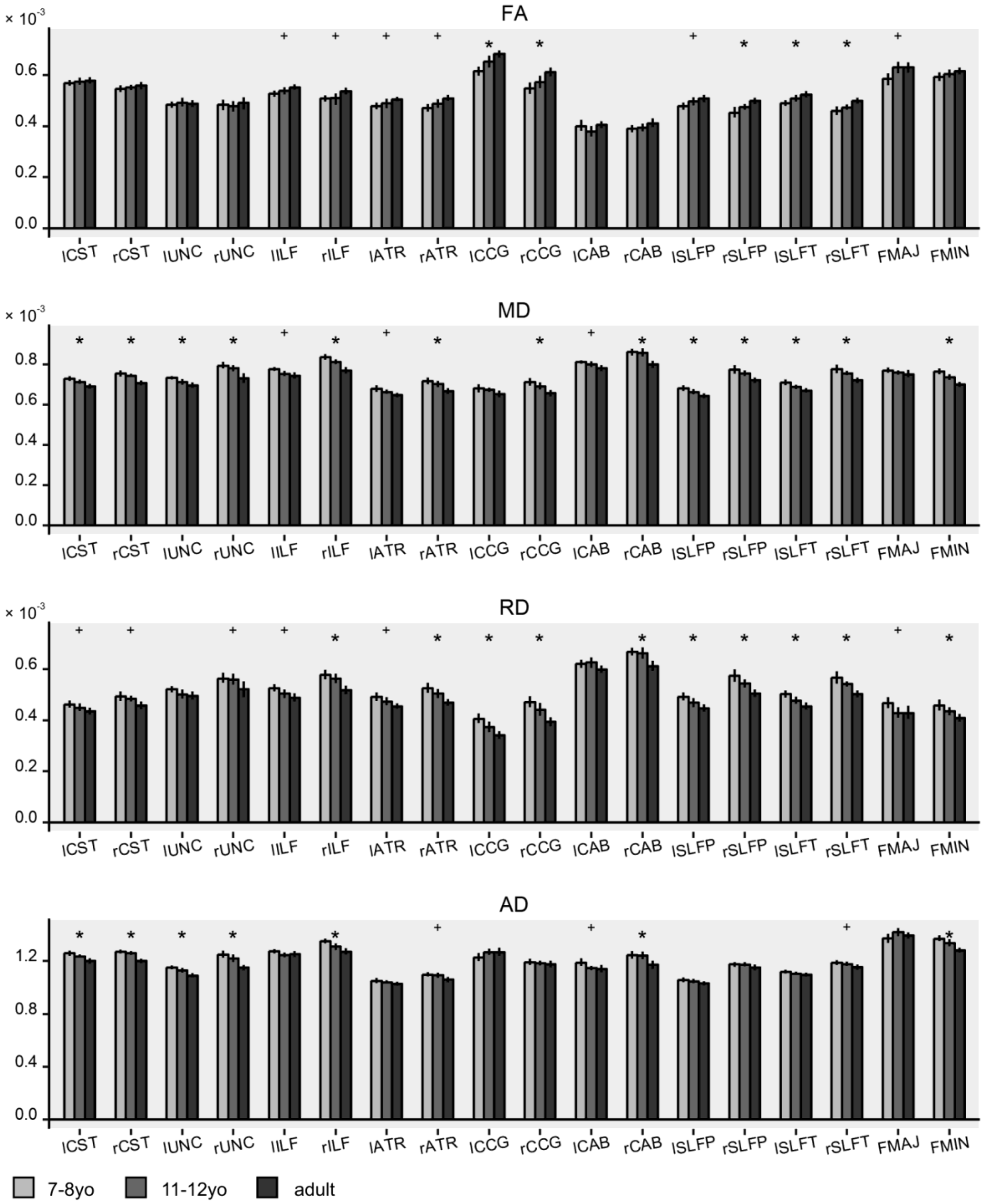

***S2 Figure**: Diffusion tensor imaging (DTI) parameters fractional anisotropy (FA), mean diffusivity (MD), radial diffusivity (RD) and axial diffusivity (AD) for 18 major white matter tracts in three age groups. Light gray bars = 7-8-year-old children, medium gray bars = 11-12-year-old children, dark gray bars = adults. Error bars show 95% confidence intervals for the mean. Asterix and plus signs indicate significance with Bonferroni correction (α = .0027) and without correction for multiple comparisons (α = .05), respectively.*

### Tables

#### S1 Table

**S1 Table**: Myelin water fraction weighted mean age effect ANOVA

| ROI | test statistic | | *p*-value | effect size |
|---|---|---|---|---|
| lCST | $F(2,45) =$ | 34.08 | $p < .00001$ | $\eta^2 = .602$ |
| rCST | $F(2,45) =$ | 40.11 | $p < .00001$ | $\eta^2 = .641$ |
| lUNC | $F(2,45) =$ | 7.90 | $p = .00114$ | $\eta^2 = .260$ |
| rUNC | $F(2,45) =$ | 5.30 | $p = .00856$ | $\eta^2 = .191$ |
| lILF | $F(2,45) =$ | 4.48 | $p = .01680$ | $\eta^2 = .166$ |
| rILF | $F(2,45) =$ | 10.45 | $p = .00019$ | $\eta^2 = .317$ |
| lATR | $F(2,45) =$ | 14.88 | $p = .00001$ | $\eta^2 = .398$ |
| rATR | $F(2,45) =$ | 9.73 | $p = .00031$ | $\eta^2 = .302$ |
| lCCG | $F(2,45) =$ | 15.33 | $p < .00001$ | $\eta^2 = .405$ |
| rCCG | $F(2,45) =$ | 7.13 | $p = .00204$ | $\eta^2 = .241$ |
| lCAB | $F(2,45) =$ | 3.928 | $p = .02680$ | $\eta^2 = .149$ |
| rCAB | $F(2,45) =$ | 3.76 | $p = .03100$ | $\eta^2 = .143$ |
| lSLFP | $F(2,45) =$ | 20.94 | $p < .00001$ | $\eta^2 = .482$ |
| rSLFP | $F(2,45) =$ | 53.53 | $p < .00001$ | $\eta^2 = .704$ |
| lSLFT | $F(2,45) =$ | 15.78 | $p < .00001$ | $\eta^2 = .412$ |
| rSLFT | $F(2,45) =$ | 41.21 | $p < .00001$ | $\eta^2 = .647$ |
| FMAJ | $F(2,45) =$ | 13.74 | $p = .00002$ | $\eta^2 = .379$ |
| FMIN | $F(2,45) =$ | 5.575 | $p = .00687$ | $\eta^2 = .199$ |

### S2 Table

**S2 Table**: Myelin water fraction weighted mean age effect planned comparison *t*-tests

| ROI | contrast | test statistic | | *p*-value | effect size |
|---|---|---|---|---|---|
| lCST | 7-8yo vs 11-12yo | $t(45)$ = | 2.75 | $p$ = .00849 | $r$ = .380 |
| | children vs adult | $t(45)$ = | 8.19 | $p$ < .00001 | $r$ = .773 |
| rCST | 7-8yo vs 11-12yo | $t(45)$ = | 2.89 | $p$ = .00584 | $r$ = .396 |
| | children vs adult | $t(45)$ = | 8.87 | $p$ < .00001 | $r$ = .797 |
| lUNC | 7-8yo vs 11-12yo | $t(45)$ = | 0.54 | $p$ = .59285 | $r$ = .080 |
| | children vs adult | $t(45)$ = | 3.76 | $p$ = .00050 | $r$ = .488 |
| rUNC | 7-8yo vs 11-12yo | $t(45)$ = | 0.43 | $p$ = .66680 | $r$ = .064 |
| | children vs adult | $t(45)$ = | 3.07 | $p$ = .00360 | $r$ = .416 |
| lILF | 7-8yo vs 11-12yo | $t(45)$ = | 1.88 | $p$ = .06700 | $r$ = .269 |
| | children vs adult | $t(45)$ = | 2.93 | $p$ = .00530 | $r$ = .400 |
| rILF | 7-8yo vs 11-12yo | $t(45)$ = | 0.94 | $p$ = .35000 | $r$ = .139 |
| | children vs adult | $t(45)$ = | 4.41 | $p$ = .00006 | $r$ = .550 |
| lATR | 7-8yo vs 11-12yo | $t(45)$ = | 2.18 | $p$ = .03500 | $r$ =. 308 |
| | children vs adult | $t(45)$ = | 5.45 | $p$ < .00001 | $r$ = .630 |
| rATR | 7-8yo vs 11-12yo | $t(45)$ = | 1.44 | $p$ = .15600 | $r$ = .210 |
| | children vs adult | $t(45)$ = | 4.37 | $p$ = .00007 | $r$ = .546 |
| lCCG | 7-8yo vs 11-12yo | $t(45)$ = | 2.32 | $p$ = .02520 | $r$ = .326 |
| | children vs adult | $t(45)$ = | 5.53 | $p$ < .00001 | $r$ = .636 |
| rCCG | 7-8yo vs 11-12yo | $t(45)$ = | 1.52 | $p$ = .13491 | $r$ = .221 |
| | children vs adult | $t(45)$ = | 3.77 | $p$ = .00047 | $r$ = .490 |
| lCAB | 7-8yo vs 11-12yo | $t(45)$ = | -0.56 | $p$ = .58080 | $r$ = .083 |
| | children vs adult | $t(45)$ = | 2.20 | $p$ = .03330 | $r$ = .311 |
| rCAB | 7-8yo vs 11-12yo | $t(45)$ = | -0.89 | $p$ = .37590 | $r$ = .132 |
| | children vs adult | $t(45)$ = | 1.90 | $p$ = .06340 | $r$ = .272 |
| lSLFP | 7-8yo vs 11-12yo | $t(45)$ = | 2.99 | $p$ = .00453 | $r$ = .407 |
| | children vs adult | $t(45)$ = | 6.47 | $p$ < .00001 | $r$ = .694 |
| rSLFP | 7-8yo vs 11-12yo | $t(45)$ = | 3.14 | $p$ = .00300 | $r$ = .424 |
| | children vs adult | $t(45)$ = | 10.21 | $p$ < .00001 | $r$ = .835 |
| lSLFT | 7-8yo vs 11-12yo | $t(45)$ = | 2.82 | $p$ = .00719 | $r$ = .387 |
| | children vs adult | $t(45)$ = | 5.61 | $p$ < .00001 | $r$ = .641 |
| rSLFT | 7-8yo vs 11-12yo | $t(45)$ = | 2.98 | $p$ = .00460 | $r$ = .406 |
| | children vs adult | $t(45)$ = | 9.00 | $p$ < .00001 | $r$ = .801 |
| FMAJ | 7-8yo vs 11-12yo | $t(45)$ = | 2.80 | $p$ = .00757 | $r$ = .385 |
| | children vs adult | $t(45)$ = | 5.22 | $p$ < .00001 | $r$ = .614 |
| FMIN | 7-8yo vs 11-12yo | $t(45)$ = | 0.49 | $p$ = .62340 | $r$ = .074 |
| | children vs adult | $t(45)$ = | 3.17 | $p$ = .00276 | $r$ = .427 |

### S3 Table

**S3 Table**: Fractional anisotropy weighted mean age effect
ANOVA

| ROI | test statistic | | *p*-value | effect size |
|---|---|---|---|---|
| lCST | $F(2,45) =$ | 0.66 | $p = .52300$ | $\eta^2 = .028$ |
| rCST | $F(2,45) =$ | 0.76 | $p = .47300$ | $\eta^2 = .033$ |
| lUNC | $F(2,45) =$ | 0.33 | $p = .71800$ | $\eta^2 = .015$ |
| rUNC | $F(2,45) =$ | 0.31 | $p = .73700$ | $\eta^2 = .013$ |
| lILF | $F(2,45) =$ | 3.96 | $p = .02600$ | $\eta^2 = .150$ |
| rILF | $F(2,45) =$ | 3.45 | $p = .04040$ | $\eta^2 = .133$ |
| lATR | $F(2,45) =$ | 3.81 | $p = .02970$ | $\eta^2 = .145$ |
| rATR | $F(2,45) =$ | 6.65 | $p = .00294$ | $\eta^2 = .228$ |
| lCCG | $F(2,45) =$ | 13.79 | $p = .00002$ | $\eta^2 = .380$ |
| rCCG | $F(2,45) =$ | 8.91 | $p = .00055$ | $\eta^2 = .284$ |
| lCAB | $F(2,45) =$ | 1.76 | $p = .18300$ | $\eta^2 = .073$ |
| rCAB | $F(2,45) =$ | 2.14 | $p = .13000$ | $\eta^2 = .087$ |
| lSLFP | $F(2,45) =$ | 4.14 | $p = .02240$ | $\eta^2 = .156$ |
| rSLFP | $F(2,45) =$ | 7.89 | $p = .00116$ | $\eta^2 = .260$ |
| lSLFT | $F(2,45) =$ | 7.72 | $p = .00132$ | $\eta^2 = .255$ |
| rSLFT | $F(2,45) =$ | 7.43 | $p = .00163$ | $\eta^2 = .248$ |
| FMAJ | $F(2,45) =$ | 5.15 | $p = .00969$ | $\eta^2 = .186$ |
| FMIN | $F(2,45) =$ | 1.97 | $p = .15200$ | $\eta^2 = .080$ |

### S5 Table

**S5 Table**: Radial diffusivity weighted mean age effect
ANOVA

| ROI | test statistic | | *p*-value | effect size |
|---|---|---|---|---|
| lCST | $F(2,45) =$ | 3.31 | $p = .04580$ | $\eta^2 = .128$ |
| rCST | $F(2,45) =$ | 4.58 | $p = .01540$ | $\eta^2 = .169$ |
| lUNC | $F(2,45) =$ | 2.94 | $p = .06320$ | $\eta^2 = .115$ |
| rUNC | $F(2,45) =$ | 3.26 | $p = .04790$ | $\eta^2 = .126$ |
| lILF | $F(2,45) =$ | 5.76 | $p = .00592$ | $\eta^2 = .204$ |
| rILF | $F(2,45) =$ | 11.93 | $p = .00007$ | $\eta^2 = .347$ |
| lATR | $F(2,45) =$ | 5.65 | $p = .00649$ | $\eta^2 = .201$ |
| rATR | $F(2,45) =$ | 10.11 | $p = .00024$ | $\eta^2 = .310$ |
| lCCG | $F(2,45) =$ | 11.89 | $p = .00007$ | $\eta^2 = .346$ |
| rCCG | $F(2,45) =$ | 12.07 | $p = .00006$ | $\eta^2 = .349$ |
| lCAB | $F(2,45) =$ | 2.49 | $p = .09430$ | $\eta^2 = .100$ |
| rCAB | $F(2,45) =$ | 10.13 | $p = .00023$ | $\eta^2 = .311$ |
| lSLFP | $F(2,45) =$ | 7.18 | $p = .00196$ | $\eta^2 = .242$ |
| rSLFP | $F(2,45) =$ | 11.33 | $p = .00010$ | $\eta^2 = .335$ |
| lSLFT | $F(2,45) =$ | 10.40 | $p = .00019$ | $\eta^2 = .316$ |
| rSLFT | $F(2,45) =$ | 11.91 | $p = .00007$ | $\eta^2 = .346$ |
| FMAJ | $F(2,45) =$ | 3.79 | $p = .03010$ | $\eta^2 = .144$ |
| FMIN | $F(2,45) =$ | 7.13 | $p = .00204$ | $\eta^2 = .241$ |

### S4 Table

**S4 Table**: Mean diffusivity weighted mean age effect
ANOVA

| ROI | test statistic | | *p*-value | effect size |
|---|---|---|---|---|
| lCST | $F(2,45) =$ | 9.39 | $p = .00039$ | $\eta^2 = .294$ |
| rCST | $F(2,45) =$ | 11.97 | $p = .00007$ | $\eta^2 = .347$ |
| lUNC | $F(2,45) =$ | 8.16 | $p = .00094$ | $\eta^2 = .266$ |
| rUNC | $F(2,45) =$ | 10.18 | $p = .00023$ | $\eta^2 = .312$ |
| lILF | $F(2,45) =$ | 6.55 | $p = .00319$ | $\eta^2 = .225$ |
| rILF | $F(2,45) =$ | 19.17 | $p < .00001$ | $\eta^2 = .460$ |
| lATR | $F(2,45) =$ | 5.45 | $p = .00758$ | $\eta^2 = .195$ |
| rATR | $F(2,45) =$ | 10.29 | $p = .00021$ | $\eta^2 = .314$ |
| lCCG | $F(2,45) =$ | 2.92 | $p = .06420$ | $\eta^2 = .115$ |
| rCCG | $F(2,45) =$ | 9.78 | $p = .00030$ | $\eta^2 = .303$ |
| lCAB | $F(2,45) =$ | 5.33 | $p = .00839$ | $\eta^2 = .191$ |
| rCAB | $F(2,45) =$ | 12.83 | $p = .00004$ | $\eta^2 = .363$ |
| lSLFP | $F(2,45) =$ | 6.88 | $p = .00247$ | $\eta^2 = .234$ |
| rSLFP | $F(2,45) =$ | 9.30 | $p = .00042$ | $\eta^2 = .293$ |
| lSLFT | $F(2,45) =$ | 9.80 | $p = .00029$ | $\eta^2 = .303$ |
| rSLFT | $F(2,45) =$ | 10.09 | $p = .00024$ | $\eta^2 = .310$ |
| FMAJ | $F(2,45) =$ | 1.59 | $p = .21500$ | $\eta^2 = .066$ |
| FMIN | $F(2,45) =$ | 19.95 | $p < .00001$ | $\eta^2 = .470$ |

### S6 Table

**S6 Table**: Axial diffusivity weighted mean age effect
ANOVA

| ROI | test statistic | | *p*-value | effect size |
|---|---|---|---|---|
| lCST | $F(2,45) =$ | 10.62 | $p = .00017$ | $\eta^2 = .321$ |
| rCST | $F(2,45) =$ | 22.03 | $p < .00001$ | $\eta^2 = .495$ |
| lUNC | $F(2,45) =$ | 12.55 | $p = .00005$ | $\eta^2 = .358$ |
| rUNC | $F(2,45) =$ | 15.84 | $p < .00001$ | $\eta^2 = .413$ |
| lILF | $F(2,45) =$ | 2.46 | $p = .09700$ | $\eta^2 = .099$ |
| rILF | $F(2,45) =$ | 11.50 | $p = .00009$ | $\eta^2 = .338$ |
| lATR | $F(2,45) =$ | 1.96 | $p = .15200$ | $\eta^2 = .080$ |
| rATR | $F(2,45) =$ | 4.86 | $p = .01230$ | $\eta^2 = .178$ |
| lCCG | $F(2,45) =$ | 2.27 | $p = .11500$ | $\eta^2 = .092$ |
| rCCG | $F(2,45) =$ | 0.40 | $p = .67100$ | $\eta^2 = .018$ |
| lCAB | $F(2,45) =$ | 3.85 | $p = .02870$ | $\eta^2 = .146$ |
| rCAB | $F(2,45) =$ | 7.06 | $p = .00215$ | $\eta^2 = .239$ |
| lSLFP | $F(2,45) =$ | 1.88 | $p = .16500$ | $\eta^2 = .077$ |
| rSLFP | $F(2,45) =$ | 1.71 | $p = .19200$ | $\eta^2 = .071$ |
| lSLFT | $F(2,45) =$ | 2.83 | $p = .06960$ | $\eta^2 = .112$ |
| rSLFT | $F(2,45) =$ | 3.35 | $p = .04390$ | $\eta^2 = .130$ |
| FMAJ | $F(2,45) =$ | 2.23 | $p = .11900$ | $\eta^2 = .090$ |
| FMIN | $F(2,45) =$ | 15.93 | $p < .00001$ | $\eta^2 = .415$ |

### Text

#### S1 Text

*Methods*

In a supplementary analysis, we investigated if DTI parameters mirrored the observed MWI effects. To this end, diffusion tensors were fit to each brain voxel in participants' individual native diffusion space with FSL DTIFIT and mean, radial, and axial diffusivity (MD, RD, AD), as well as fractional anisotropy (FA) were computed. Analogous to MWF, for each tract, we calculated weighted mean DTI parameters.

*Results*

DTI parameters revealed a mixed pattern in relation to the MWF age effect. In any given tract, we defined a diverging result as a bonferroni-corrected significant age effect for MWF but no indication of significance at all, i.e. $\alpha > .05$. FA, the DTI parameter most-widely interpreted as "structural integrity", increased with age in five tracts only (lCCG, rCCG, rSLFP, lSLFT, rSLFT, range of $\eta^2$ = [.248 .380], S2 Figure, S3 Table). In another six tracts, FA increased, but did not meet our significance threshold (lILF, rILF, lATR, rATR, lSLFP, FMAJ, range of $\eta^2$ = [.132 .228]). Critically, in three tracts (lCST, rCST, lUNC) that showed stark MWF increases with age, FA did not show effects of age and featured near-to-zero effect sizes ($\eta^2$ = .028, .033, and .015). MD decreased with age in 13 out of 18 tracts and showed non-significant decreases in another three tracts (lILF, lATR, lCAB, Figure 2, S4 Table). Strikingly, in the lCCG and FMAJ, we did not find any age group differences for MD ($\eta^2$ = .114 and .066), while these tracts showed clear MWF increases. RD decreased with age in 10 out of 18 tracts and showed non-significant decreases in another six tracts (lCST, rCST, rUNC, lILF, lATR, FMAJ, Figure 2, S5 Table). In the lUNC and FMAJ, we did not find RD age group differences ($\eta^2$ = .115 and .144), despite strong MWF differences. AD decreased with age in seven out of 18 tracts and showed non-significant decreases in another three tracts (rATR, lCAB, rSLFT, Figure 2, S6 Table). Thus, AD did not reveal any effect of age in seven tracts that featured significant effects of age for MWF (lATR, lCCG, rCCG, lSLFP, rSLFP, lSLFT, FMAJ).

In summary, we found that while age group differences in MD and RD mirror MWF differences in great proportion of tracts, FA or AD age group differences correspond to MWF results only in few tracts.

*Influence of tract size on DTI parameters*

DTI metrics (especially FA), could be influenced by tract volume, as smaller tracts have a larger border-to-volume ratio. Complex fiber architecture that decreases FA but does not influence functioning or myelination should be more common at border regions of a tract (Lebel and Deoni,

2018). Moreover, partial volume effects, i.e. voxels bordering white and gray matter and thus show biased metrics are more common in small volume tracts. When comparing tract volume between age groups, we only found significant differences for the CST (left: $F(2,45) = 7.00$, $p = .00225$, $\eta^2 = .237$; right: $F(2,45) = 15.85$, $p = .00001$, $\eta^2 = .413$). Strikingly, the CST exhibited the most robust MWF increase, but no FA increase. Note that the influence of bordering regions in our data should be low, as bordering regions should have a low *a posteriori* tractography probability and contribute less to the weighted mean. Thus, while lacking FA age effects in the CST might be explained by tract size differences between age groups, for all other tracts, volume seems an unlikely bias.

#### S2 Text

*Discussion*

Our multimethod approach of combining MWI with DTI enables us to relate our MWF findings to widely used DTI parameters. We demonstrate that interpreting DTI parameters as a proxy for myelination is less accurate than MWI in detecting age differences. Still, MD and RD can reproduce myelin-specific age effects in most tracts. This agrees with previous findings of high sensitivity of RD to myelination in histological animal samples (Song et al., 2005; Song et al., 2003; Song et al., 2002) and a high correlation of RD and MWF in late childhood (Geeraert et al., 2018). In contrast, FA failed to reproduce myelin-specific age effects in many tracts. This leads us to endorse previous notes of caution against interpreting FA as a marker for myelination (e.g. Jones et al., 2013) and suggests that developmental DTI studies that commonly found FA increases alongside MD and RD decreases (for an overview, see Lebel and Deoni, 2018) might have tapped distinct processes. AD also overwhelmingly failed to reproduce myelin-specific age effects. This agrees with the inconclusive picture of mixed results of AD development in childhood, suggesting no robust age effects of AD (for an overview, see Lebel and Deoni, 2018), as well as with histological animal studies indicating that AD is not sensitive to myelination (Song et al., 2002), but rather to axonal degradation (Song et al., 2003). Thus, AD does not hold as a useful marker of myelin development from childhood to adulthood.

### Supplementary material references

Geeraert, B.L., Lebel, R.M., Mah, A.C., Deoni, S.C., Alsop, D.C., Varma, G., Lebel, C., 2018. A comparison of inhomogeneous magnetization transfer, myelin volume fraction, and diffusion tensor imaging measures in healthy children. Neuroimage 182, 343–350. 10.1016/j.neuroimage.2017.09.019.

Jones, D.K., Knösche, T.R., Turner, R., 2013. White matter integrity, fiber count, and other fallacies: the do's and don'ts of diffusion MRI. Neuroimage 73, 239–254. 10.1016/j.neuroimage.2012.06.081.

Lebel, C., Deoni, S., 2018. The development of brain white matter microstructure. Neuroimage 182, 207–218. 10.1016/j.neuroimage.2017.12.097.

Song, S.-K., Sun, S.-W., Ju, W.-K., Lin, S.-J., Cross, A.H., Neufeld, A.H., 2003. Diffusion tensor imaging detects and differentiates axon and myelin degeneration in mouse optic nerve after retinal ischemia. Neuroimage 20 (3), 1714–1722. 10.1016/j.neuroimage.2003.07.005.

Song, S.-K., Sun, S.-W., Ramsbottom, M.J., Chang, C., Russell, J., Cross, A.H., 2002. Dysmyelination Revealed through MRI as Increased Radial (but Unchanged Axial) Diffusion of Water. Neuroimage 17 (3), 1429–1436. 10.1006/nimg.2002.1267.

Song, S.-K., Yoshino, J., Le, T.Q., Lin, S.-J., Sun, S.-W., Cross, A.H., Armstrong, R.C., 2005. Demyelination increases radial diffusivity in corpus callosum of mouse brain. Neuroimage 26 (1), 132–140. 10.1016/j.neuroimage.2005.01.028.
